# A novel maxillofacial technique to deliver drugs to the visual pathway, the optic nerve and the retina using the glymphatic system

**DOI:** 10.1101/2022.03.23.485523

**Authors:** Jayamuruga Pandian Arunachalam, Rahini Rajendran, Subbulakshmi Chidambaram, Srikanth Krishnagopal, S. Bhavani, V. Subha, Veeran Veeravarmal, Harikrishnan Prasad, U.R Anoop, Kavita Verma

**Affiliations:** Central Inter-Disciplinary Research Facility, Sri Balaji Vidyapeeth [Deemed to be University], SBV-Mahatma Gandhi Medical College & Research Institute campus, Pondicherry 607402, India; Sensory Neural Engineering and Cell Therapeutics lab Department of Biochemistry and Molecular Biology Pondicherry Central University, Puducherry 605014, India; Professor and Head Department of Ophthalmology, Sri Balaji Vidyapeeth [Deemed to be University] Mahatma Gandhi Medical College & Research Institute, Pondicherry 607402, India; Department of Pathology, Sri Balaji Vidyapeeth [Deemed to be University] Mahatma Gandhi Medical College & Research Institute, Pondicherry 607402, India; Department of Pharmacology, Sri Balaji Vidyapeeth [Deemed to be University] Mahatma Gandhi Medical College & Research Institute, Pondicherry 607402, India; Department of Oral Pathology and Microbiology, Cuddalore Government Dental College and Hospital, Annamalai Nagar, Chidhambaram, Tamilnadu, India; Department of Oral Pathology and Microbiology, KSR Institute of Dental Science and Research, Tiruchengode, Tamilnadu, India; UR Anoop Research Group, Pondicherry, India. 605008.

**Keywords:** Retina, Optic nerve, Topical, Subconjunctival. Intravitreal, Maxillofacial, glymphatic pathway.

## Abstract

The blood neural barriers are formidable barriers that prevent delivery of drugs into the eye and the brain. Currently, large biologics are used for treating many diseases of the eye and the brain. But the blood retinal barrier and the blood brain barrier prevent entry of these useful drugs into the eye and the brain respectively. Hence, there is a need for a drug delivery technique to bypass these natural barriers.

Bevacizumab (Avastin) is a full length humanized monoclonal antibody with a molecular weight of 149kda. It binds to circulating vascular endothelial growth factor A and prevents interaction with vascular endothelial growth receptors. This results in blocking endothelial response and tumor vascularization. Bevacizumab is used to prevent choroidal neovascularization in age related macular degeneration and to prevent retinal neovascularization in diabetic retinopathy. It is also used for treating certain types of brain cancer. But it crosses neither the blood retinal barrier nor the blood brain barrier.

This in vivo mouse study assessed the delivery of a commercial formulation of bevacizumab into the retina of mice using different ocular techniques like topical drops, subconjunctival injection, intravitreal injection and a novel maxillofacial technique. The objective was to assess whether the novel maxillofacial drug delivery technique could deliver drugs into the retina and to compare the efficacy of the maxillofacial technique with the mentioned ocular techniques. The distribution of bevacizumab was further compared with retinoschisin which is a small protein with a molecular weight of 24kda.

Intravitreal injection and maxillofacial technique were effective in delivering drugs into the retina. In addition, the maxillofacial technique could target the brain including regions involved in the visual pathway and the optic nerve. The glymphatic pathway could also be targeted for drug delivery. Drug was also detected in the contralateral optic nerve and retina.

Based on our study findings, we propose a new concept to explain the presence of the drug in the contralateral eye. We propose that after maxillofacial drug delivery, early distribution of the drug can occur in the CSF at the optic chiasma from the brain via the glymphatic system. In the case of the intravitreal injection, the drug from the experimental eye may be cleared through the glymphatic pathway of the ipsilateral optic nerve into the CSF surrounding the nerve. The crossover of the ipsilateral optic nerve fibers at the optic chiasma can result in further distribution of the drug in the CSF at the optic chiasma. From the region of the optic chiasma, the drug can distribute into the CSF surrounding the contralateral optic nerve. The drug can be then distributed into the contralateral optic nerve through the glymphatic pathway and be delivered into the contralateral eye. The drug can be further cleared into the aqueous of the contralateral eye through the anterior clearance pathway.

## INTRODUCTION

The ocular barriers namely the cornea, the blood aqueous barrier, the inner retinal barrier and the outer retinal barrier of the eye pose a challenge to ocular drug delivery. Therefore, different ocular drug delivery techniques are used for delivering drugs into the eye^1^.Topical and systemic routes require high doses of drugs to obtain the therapeutic range in the eye. Hence, locally invasive techniques like sub-conjunctival and intra-vitreous are used to deliver drugs into the posterior compartment. As the elimination rate of the injected drugs is fast, multiple injections are required. This can lead to complications and lack of patient compliance^1^. Therefore, a drug delivery system that can target the retina and the optic nerve effectively and non-invasively holds immense potential.

Similarly, the blood brain barrier prevents the entry of large biologics into the brain. Therefore, useful antibodies like bevacizumab which is FDA approved for brain cancer cannot enter the brain^2^. Many techniques are used to circumvent the blood brain barrier. The strategy to cross the blood brain barrier by increasing the plasma drug concentrations with high dose intravenous infusions, bolus injections and intra-arterial delivery resulted in systemic complications. Invasive techniques like convection enhanced delivery, intra-cerebroventricular infusion, intra-cerebral injection, polymer implants placed directly into the brain and techniques that open the blood brain barrier using osmotic disruption, bradykinin analogs and ultrasound make the brain prone to infection. Non-invasive techniques like the pharmacological approaches, the physiological approaches and nasal drug delivery do not provide controlled and continuous drug delivery ^3^.

Many neurodegenerative conditions involving the brain also involve the eye^4^. Hence, a drug delivery technique that can target both the brain and the eye can be useful in many age-related and neurodegenerative conditions. This in vivo mouse study assessed whether drugs could be delivered into the retina through the glymphatic pathway of the optic nerve using a novel maxillofacial drug delivery technique. Maxillofacial drug delivery was further compared with intravitreal injection, subconjunctival injection and topical eye drops.

## MATERIALS & METHOD

### 2.1 Animals and protein administration

Wild type male C57BL/6J mice (procured from Biogen Laboratory animal facility, Bangalore) of approximately 8-9 months old with an average weight of 35 grams were used for the study. The animals were anesthetized with Ketamine (Neon Laboratories Ltd) 80-120mg/Kg weight of mice and Xylazine (Indian Immunological Ltd) 7-10mg/Kg weight of mice. Povidone-Iodine 5% was applied to both eyes. 250µg of a commercial preparation of Bevacizumab (Avastin) (Roche Chemicals Ltd) and 10ug recombinant RS1-FLAG protein was used in the study. The drug was administered through either topical, subconjunctival, intravitreal or maxillofacial routes. In the topical group, eye drops were used in the right eye. In the sub-conjunctival group and the intravitreal group, the right eyes were injected with Bevacizumab or RS1-FLAG using a 30-gauge needle. The left eye received no injections. In the control, 1XPBS was injected into the animal’s right eye. Eyes were monitored closely for any signs of adverse effects. All the ocular injections were done by experienced ophthalmologist.

In the maxillofacial group, the drug was delivered to the connective tissue of the respiratory lining mucosa of either the right or the left maxillary sinus using a novel implantable drug delivery system. The implant was inserted in the maxillary sinus region by making a midline incision in the dorsum of the nose.

Following administration of the drug, the animals were perfused (perfused by an experienced animal surgeon) using 10% formalin after 24 hours or after 48hours. The brain, the right, the left eye, the liver and the kidneys were collected for immunohistochemical analysis.

For LCMS/MS and western blot analysis, 1ml of blood was collected from the animals. Then the animals were perfused with saline. The brain, the right eye, the left eye, the liver and the kidneys were collected and the total proteins were isolated from the tissues using 1XRIPA buffer (25mM Tris-Cl (pH 7.6), 150mM NaCl, 1% Triton X-100, 0.1% SDS, and 0.5% Sodium deoxycholate). The isolated proteins were quantified using Bradford assay and were aliquoted and stored at -80°C for further analysis.

### 2.2 LCMS/MS analysis

Trypsin digested protein samples were analyzed by Impact HD Q-TOF (Bruker, Germany) mass spectrometer. We have selected specific peptides related to the standards (Avastin and RS1-His tag) by simulating standard sequence in sequence editor using trypsin enzyme. The presence of those peptides was checked in trypsin digested standard samples. Based on the data three peptides were selected for Avastin (two for heavy chain, one for light chain) and two peptides were selected for RS1.

Avastin peptides eluted at 15.6 min (light chain - mol wt 1877.9112 da DIQMTQSPSSLSASVGDR), 19.3min (heavy chain - mol wt 1185.6318da GPSVFPLAPSSK) and 22.3min (mol wt 1806.9889 da VVSVLTVLHQDWLNGK)

RS1 peptide eluted at 18.9min (mol wt 520.26da MSRK) and 19.4min (mol wt 1693.984 da TSTVQNLLRPPIISR)

Those mol wt and retention time were checked for all the trypsin digested samples and the area was tabulated. Overnight trypsin digestion was carried out, Zorbax Eclipse plus C18 (4.6X100mm,3.5µ) column was used for separation, acetonitrile and 0.1% formic acid as a mobile phase at the flow rate 0.3ml/min.

#### Results

The control samples were negative for Bevacizumab. All the samples from the study group were positive for Bevacizumab.

### 2.3 Immunohistochemistry

For immunohistochemical analysis, 10% formalin was used to fix the tissues. After processing and embedding the tissues, thin sections (3.5-5µm) were made on the IHC slides. The sections were deparaffinized at 60°C in a hot air oven for1h . Then, the sections were washed two times with xylene for 20min each and rehydrated with decreasing concentrations of ethanol (90% to 50%). The sections were washed with distilled water for 5min. Antigen retrieval was done by immersing the slides in boiling Tris EDTA buffer (pH 9) for 40min. The slides with the sections were allowed to cool and washed with wash buffer for 5min. Then the slides were incubated with 3% H2O2 for 10min. After incubation, the slides were washed two times with a wash buffer for 5min each. 5% Bovine Serum Albumin (BSA) was used as blocking agent and was incubated for 15min at 37°C. The slides were incubated with the primary antibody Anti-FLAG rabbit mAb [CST #14793] (1:800 dilution) for RS1-FLAG treated animal tissues for 1h at 37°C in a humidified chamber. The slides were washed twice with a wash buffer for 5min each. Then the secondary antibody [Agilent Dako Envision^s^ FLEX HRP #SM802] was added and incubated for 30min at 37°C in a humidified chamber. After incubation, the slides were washed, and a freshly prepared DAB substrate mixture was applied to the slides and incubated for 10min in dark, and the slides were washed twice with wash buffer for 5min each. Enough volume of Mayer’s hematoxylin was applied over the sections and incubated for 30sec, and then the slides were washed with distilled water for 5min and rinsed with tap water. The sections were dehydrated by dipping the slides in 100% Ethanol and mounted with a coverslip using a mounting medium. For Avastin-treated animals, after blocking, direct secondary antibody [CST #32935 Anti Human IgG HRP 1:1000] was added to detect the anti-VEGF signals. Suitable positive and negative controls were used for result interpretation. For positive control #CST 2056 PKCα Rabbit Ab (1:100) was used.

#### RESULTS

**a) Topical route:**

24 hours after topical application of Avastin as eye drops, no staining was detected in the retina. The lumen of blood vessels in the inner retina showed staining probably due to systemic distribution of the drug. 48 hours after the topical application of Avastin, no staining was detected in the retina. Retrobulbar tissue and the lumen of vascular channels in the optic nerve showed staining probably due to systemic distribution of the drug.

24 hours after topical application of Retinoschisin, the outer segment of the photoreceptors showed positive staining. Occasional staining was also detected in some areas of the inner nuclear layer and the ganglion cell layer. 48 hours after topical application of Retinoschisin, no staining could be detected in the outer segment of the photoreceptor layer. Lumen of a few blood vessels in the inner retinal layer showed some staining.

**b) Subconjunctival:**

24 hours after injection of Avastin, staining was detected in the outer segment of the photoreceptors, Occasional staining was seen in the inner nuclear layer and ganglion layer. The periphery of the optic nerve also showed staining. 48 hours after injection of Avastin, no staining was detected in the outer segment of the photoreceptors. Staining was detected within the optic nerve and at the periphery of the nerve. 24 hours after injection of Retinoschisin, no staining was detected in the retina. Similarly, 48 hours after injection of Retinoschisin, no staining was detected in the retina.

**c) Intravitreal: (Fig:1)**

24 hours after injection of Avastin, significant staining was detected in the outer segment of the photoreceptors. The periphery of the optic nerve also showed staining. 48 hours after injection of Avastin, the outer segment of the photoreceptors showed minimal staining. But staining was detected within the optic nerve and at the periphery of the nerve. 24 hours after injection of Retinoschisin, no staining was detected in the retina.

**d) Maxillofacial (Fig: 1,2)**

In one animal, the drug delivery system made of surgical grade stainless steel was implanted in the maxillary sinus region through an incision made in the overlying skin under anesthesia. After implantation of the device and suturing of the overlying skin, the animal was given post operative analgesics and observed for a week. The animal had normal feeding habits and the healing was uneventful. After a week, 250µg of Avastin was administered in a controlled manner through the device under anesthesia. The animal was sacrificed after 24 hours. The saline perfused tissues were collected and stored. In other animals, drug was administered immediately after implantation of the device.

24 hours after administration of Avastin, staining was detected in the outer segment of the photoreceptors in the ipsilateral eye and in the contralateral eye. Similarly staining was detected in the ipsilateral and the contralateral optic nerve. The intensity of staining was slightly more in the contralateral eye. At 48 hours after administration of Avastin, staining was lesser in the outer segment of the photoreceptor layer of both the ipsilateral and contralateral eye. Staining was present in the ipsilateral and contralateral optic nerve also. The intensity of the staining was mildly more in the contralateral eye. Staining was also detected in the superficial and deep portions of the brain. The cortex, cerebellum, the brainstem especially the pons also showed staining. Staining was also seen in the entorhinal cortex The choroid plexus also showed staining.

In the 48 hours tissue samples in which two doses were given at 0 hours and 24 hours, staining was again detected in the outer segment of the photoreceptor layer of the ipsilateral eye and contralateral eye. Staining was also detected to a lesser extent in the inner nuclear layer and the ganglion cell layer. The optic nerves also showed staining within the nerves. The staining in the contralateral optic nerve was significant. In the brain, staining was reduced though it was present in the cortex, the cerebellum and the brainstem especially at the pons. The walls of the pial vessels showed staining.

24 hours after administration of Retinoschisin, staining was detected in the outer segments of the photoreceptor layer of the ipsilateral and contralateral eyes. The contralateral optic nerve showed significant staining. 48 hours after administration of Retinoschisin, the outer segment of the photoreceptor layers of the contralateral eye also showed positive staining. Mild staining of the nerve fiber layer was also seen.

### 2.4 SDS-PAGE & Western blot analysis

12% resolving gel and 5% stacking gel were used for separating the proteins. The protein samples were prepared by boiling in 2X Laemmli buffer for 5 min at 95°C. The proteins were transferred into a 0.2µm PVDF membrane and blocked with 5% non-fat dry milk for 1h at room temperature..

For Avastin detection, direct secondary antibody (1:5000 in 5% non-fat dry milk) [CST #32935 Anti Human IgG HRP] was used.

## RESULTS

In samples collected after 24 hours and 48 hours following topical application, the drug was not detected in either of the eyes. The kidney and plasma were positive. The brain and liver were negative.

In samples collected 24 hours after subconjunctival application, the right eye, kidney and plasma were positive. The left eye, brain and liver were negative.

24 hours after the intravitreal injection, the drug was detected in the right and left eyes, liver, kidney and plasma. The brain was negative. 48 hours after the intravitreal injection, the drug was detected only in the right eye. The left eye, brain and liver were negative. The kidney and plasma were positive.

In the samples collected 24 hours after maxillofacial drug delivery, both the eyes were positive. The right and left cortex, the right and left halves of pons and the right and left halves of cerebellum were positive. The drug was also detected in liver, kidney and plasma.

## DISCUSSION

The posterior segment of the eye comprises the back two-thirds of the eye including the vitreous, the retina, the choroid and the optic nerve. ^5^. Many diseases that affect the posterior segment result in impaired vision or blindness ^6^. But delivering drugs to the posterior segment is a challenge because of the anatomic barriers and the physiologic clearance mechanisms^7^.The static barriers include the cornea, the sclera, the retina, the blood - retinal barrier and the blood - aqueous barrier, The dynamic barriers include the choroidal blood flow and the conjunctival blood flow, lymphatic clearance and tear dilution ^8^.

Various routes for administration of drugs into the retina have been investigated. These include topical, systemic, intra-ocular (intravitreal), periocular (sub-conjunctival, sub-tenon’s, posterior juxtascleral, peribulbar and retrobulbar injections), sub-retinal and suprachoroidal ^6, 7^

In this mouse study, we compared topical eye drops, sub-conjunctival injection, intravitreal injection and a novel oral and maxillofacial technique for retinal drug delivery of bevacizumab a large antibody. The distribution of bevacizumab was compared with retinoschisin, a small molecule.

(Bevacizumab) is a recombinant humanized full monoclonal antibody with a Fc fraction and has affinity for all forms of human vascular endothelial factor -A (VEGF-A). Bevacizumab has a molecular weight of 149kda and is negatively charged under physiologic conditions^9^

Intravitreal injection is currently the standard procedure in clinical practice for retinal drug delivery^5, 6^. But static barriers like the vitreous and the retina and dynamic barriers like the anterior elimination pathway via bulk aqueous flow and the posterior elimination pathway via the vitreoretinochoroidal bulk flow due to hydrostatic and osmotic pressure gradients play an important role in drug distribution and clearance^7^.

Diffusion through the vitreous is based on the physiochemical properties of the drug and the retention effect of the vitreous. Vitreous contains hyaluronic acid and collagen which provide the consistency and attract water. Hyaluronic acid is negatively charged and restricts movement of positively charged drugs. As bevacizumab is negatively charged, it can diffuse through the vitreous. The vitreous contains fewer proteins than plasma. The main proteins are collagen, albumin and immunoglobulins. Binding of drugs to the proteins can impede the transport of drugs in the vitreous. The transport of Bevacizumab is not affected by proteins^9^.

The concentration gradient of the drug at the injection site provides the force for the drug to diffuse till the concentration balance is achieved. Low molecular weight drugs diffuse faster than high molecular weight drugs. To a minor extent, convection through the vitreous takes place due to difference in pressure and temperature between the anterior chamber and the retinal surface^5^.

Elimination of the drugs from vitreous is either by metabolism or by disposal into the systemic circulation. Bevacizumab is not degraded in the eye and is cleared into the systemic circulation. Larger hydrophilic molecules are removed through the anterior segment by the aqueous humor outflow into the trabecular and uveoscleral pathways. The small lipophilic drugs permeate through the retina and the blood retinal barriers and are cleared in the posterior segment by the choroidal blood flow^5^. As bevacizumab is a large heavy drug of 149kda, the clearance is slow and therefore it has a longer half-life. Elimination of bevacizumab is considered to occur mostly through the anterior route into the systemic circulation^9^. It has been shown that intravitreal bevacizumab penetrates the anterior chamber angle, iris and ciliary body in primate studies^10^.

The retina is a cellular central nervous system tissue with 15-20 nm wide intercellular spaces that do not contain tight junctions. Both hydrophilic and lipophilic small drugs can easily permeate through the retina when compared to large drugs^7, 11^. Bevacizumab has been shown to penetrate through the retinal layers in many animal models and be cleared into the systemic circulation through the posterior route^12–16^. The Fc portion of Bevacizumab is associated with FcRn-mediated recycling within endothelial cells^17–20^. The Fc containing molecules are protected from intracellular degradation within endosomes. This decreases the systemic clearance of the drug. Bevacizumab shows a high systemic exposure when compared to other Anti-VEGF drugs^17, 18, 21^.

The intravitreal route can be used to inject large anti-VEGF protein drugs like bevacizumab directly into the vitreous to achieve very high bioavailability in the posterior segment without systemic toxicity. As the posterior transretinal and the anterior aqueous humor pathways clear injected drugs, the drug concentrations can fall below therapeutic levels over time^7^. As the large anti-VEGF drugs have long intravitreal half-life, are tolerated at higher doses and are potent even at lower concentrations, the injections can be repeated at longer intervals. But as smaller drugs are cleared at a fast rate, frequent injections may be required^6^. Therefore, patient compliance is low because of the need for repeated injections. Complications also involve vitreous hemorrhage, retinal detachment and endopthalmitis^7^.

In a study using wild type C57BL/6 mice, bevacizumab immunoreactivity could be detected even in the deeper layers of the retina half an hour after intravitreal injection. The staining was present 1 day after the injection. It started fading gradually. Strong immunoreactivity was found in the photoreceptor layer^12^.

In a laser scanning confocal study using the eyes of albino rabbits injected with 2.5 mg of Avastin 24 hours before being killed, immunoreactivity was found throughout the retina but not within Retinal Pigmented Epithelium (RPE) or choroid. Immunoreactivity was present in the Internal Limiting Membrane (ILM), the ganglion cell, the inner nuclear layer as well as the inner and outer segment layers of the photoreceptors. Most of the immunoreactivity for bevacizumab was extracellular. By day 7, there was no staining of the outer segment of the photoreceptors. Mueller cells showed reactivity. By 4 weeks no staining was detected^13^.

In another study using Dutch-belted rabbits, Avastin concentrations in the aqueous, vitreous and serum were measured after an intravitreal injection of 1.25mg of Avastin. In the injected eye, a peak concentration of 400µg/ml was achieved in the vitreous 1 day after intravitreal injection of 1.25 mg of Bevacizumab. It declined in a monoexponential fashion with a half-life of 4.32 days. In the aqueous, a peak concentration of 37.7 µg/ml was detected 3 days after the injection. In the serum, a maximum concentration of 3.3µg/ml was detected 8 days after the injection and the concentration fell below 1µg/ml 29 days after the injection^14^.

There are limitations for the rodent model when compared with the human eye. The mice and rabbit models do not have a macula or a fovea^12^. The lens is larger than humans. The vitreous volume is less. The volume of distribution is different from humans. In the rabbit, the retina is less vascular than in humans^14^.

In a study using cynomolgus monkeys, the penetration of intravitreally injected Avastin through the retina was assessed. Immunoreactivity was first detectable in the inner retinal layers and then spread to the outer layers of the retina. The initially strong staining of the inner retinal layers disappeared after a week. By day 7, the immunoreactivity was seen predominantly at the outer segments of the photoreceptor layer. It was also detectable in the RPE. By day 14, only the outer segment of the photoreceptor layer and the ILM showed staining. The choroid was stained at all the four investigated time points indicating a substantial transfer of Bevacizumab into the blood circulation The time dependency of bevacizumab distribution shows that it spreads from the inner retina to the outer segment of the photoreceptor layer where it stays for a longer period.^15^.

Intravitreal injection of bevacizumab labeled with^125^ I showed radioactive labeling in all layers of the retina including the RPE and the choroid on day 7 after the injection. Radioactivity was also concentrated in vessel walls. 1 day after the injection, radioactivity was detectable in the serum samples which was equal to 4% of the totally injected amount of radioactive protein. On day 4 and day 7, the percentage increased to 5%^15^.

In an electron microscopic study in Cynomolgus monkeys after an intravitreal injection of 1.25 mg of Avastin, a significant reduction of choriocapillaris endothelial cell fenestrations was seen as early as 24 hours indicating that the drug reached the choriocapillaris within the given period. The number of endothelial cell fenestrations decreased moderately from day 1 to day 4 indicating further penetration of the drug in the choriocapillaris. From day 7 to day 14, the number of fenestrations increased again, signifying the diminishing effects of the drug. However, the number of fenestrations were still significantly reduced when compared with control.^16^.

In our intravitreal study, the smaller retinoschisin was cleared faster than the larger bevacizumab. Bevacizumab staining was detected in the outer segment of the photoreceptors and in the optic nerve. The staining of the outer segment of the photoreceptor layer was more at 24 hours while the optic nerve showed more staining at 48 hours suggesting an additional elimination route through the optic nerve in addition to the anterior and choroidal pathways.

Following intravitreal injection, Anti-VEGF drugs have been detected in the systemic circulation. Though the quantity of the drugs that is cleared from the eye into the systemic circulation is low, suppression of systemic VEGF occurs as these drugs are potent even at subnanomolar range. A clinical prospective study on 56 patients with wet age-related macular degeneration compared the systemic exposure and systemic vascular endothelial growth factor suppression of ranibizumab, bevacizumab and aflibercept. Serum pharmacokinetics and plasma free VEGF were evaluated after the first and third intravitreal injection. Among the three drugs systemic exposure of bevacizumab was the highest and ranibizumab was the lowest. Bevacizumab was cleared relatively slowly from systemic circulation and appeared to accumulate with repeated dosing. It also reduced free VEGF in the plasma. The systemic presence of bevacizumab and the associated reduction in plasma free VEGF was very rapid and was seen at 3 hours post-treatment probably facilitated by the Fc portion^17^. A prospective, open label, non-randomized clinical trial using intravitreal Anti VEGF in patients with neo-vascular age-related macular degeneration, diabetic macular edema or retinal vein occlusion also showed that systemic exposure was the highest for Bevacizumab^18^.

A non-human primate study using positron emission tomography-computed tomography in owl monkeys for assessing systemic distribution and intravitreal pharmacokinetic properties of I-124- labeled Bevacizumab, Ranibizumab and Aflibercept found that bevacizumab had the longest intravitreal retention time. Radioactivity emission levels in collected blood samples measured by a gamma counter showed higher levels of Bevacizumab in the serum. None of three drugs accumulated in the central nervous system. In non-injected fellow eyes, all the three drugs were visible on days 1, 2 and 4. Bevacizumab was significantly high only on day 8. All the three drugs had very low levels of detection after day 8. A significantly level of bevacizumab was seen in the spleen, kidney, urinary bladder, liver, heart and distal bones.^22^.

Our western blot study at 24 hours showed signals in the injected eye, non-injected fellow eye and plasma. The brain showed no signals. Liver and kidney also showed signals. But at 48 hours, only the injected eye and plasma showed the signals. The brain did not show any signals. The kidney showed positive signals.

Subconjunctival drug delivery is a form of periocular drug delivery wherein the drug is injected under the conjunctiva that covers the sclera. The rate limiting corneal - conjunctival epithelial barrier can be bypassed and the drug can be transported through the transscleral route^5^.

Retinal bioavailability in this route is low because of the static and dynamic barriers that impede the distribution of the drug^6^. The deposited drug has to pass through static barriers like the sclera, choroid, Bruch’s membrane and RPE to reach the vitreous and the retina. This results in a steep drug concentration with the highest concentration in the sclera and the lowest at the vitreous and the retina^7^.

The dynamic barriers include the subconjunctival – episcleral blood flow and the lymphatic circulation and the choroidal circulation^8^. The robust blood and lymphatic clearance mechanism in the subconjunctival space clears the drugs fast into the systemic circulation and affects the retention time of the drug. This makes the route less effective in attaining high peak concentration at the retina and the vitreous^7^. The permeation through the sclera is size dependent. Smaller molecules pass through the sclera easily while larger molecules permeate slowly. Blood flow in the choroid is extremely high^8^. Therefore, the drugs that reach the underlying choroid are cleared fast by the choroidal blood flow into the systemic circulation. The smaller drugs are cleared at a faster rate than the larger drugs. Therefore between 80-95% of the small drugs drain into the systemic circulation. Approximately only 10% reaches the aqueous humor and only 0.1% reaches the retina^5^.

The procedure is less invasive and higher volumes of the drug can be given when compared with the intravitreal route. Enzymatic degradation of the drug is also low due to lack of enzymes at the injection site. The drug delivery to the back of the eye is better when compared with topical and systemic routes^5^. The risks of endophthalmitis and cataract can be avoided by this route.

In our IHC study, after subconjunctival injection, limited bevacizumab was detected at 24 hours in the outer segment of the photoreceptor layer and at the periphery of the optic nerve. At 48 hours staining was not detected in the outer segment of the photoreceptor layer. The optic nerve showed staining suggestive of posterior clearance through the optic nerve. But retinoschisin was not detected in the retina probably because of the faster clearance as it is a small molecule.

Our western blot study further detected signals in the injected eye, kidney and plasma. The brain, non-injected fellow eye and the liver did not show any signals.

Topical drug delivery is the most common route used for ocular disease. But the anatomical and physiological barriers prevent topical drugs from reaching the posterior segment. The static anatomic barriers are the corneal epithelium, conjunctival epithelium, sclera, choroid, Bruch’s membrane, RPE and the retina. The conjunctival epithelium is more permeable than the corneal epithelium because of the leaky goblet cells and the larger surface area. The dynamic barrier includes tear turnover which results in washing away of the drug from the ocular surface into the nasolacrimal duct within an average of five minutes. The sub-conjunctival -episcleral blood and lymphatic flow and the choriocapillaris blood flow further clear the drug that penetrates into the underlying tissues. Though permeation barriers to topical drug delivery are similar to sub- conjunctival injection, the conjunctival epithelium further decreases the permeation by approximately fivefold. Some studies have shown that topically administered drugs do reach the back of the eye. But the intravitreal and intraretinal bioavailability is very low^7^.

In a study, 1% bevacizumab was applied topically to the cornea of a group of BALB/c mice and a single dose of 0.5 mg Bevacizumab was injected subconjunctivally in another group. The corneal penetration of topical and subconjunctival Bevacizumab was assessed. Bevacizumab is a full-length immunoglobulin with a 12 nm long, Y-shaped configuration Its three arms are approximately 3.5 nm in diameter. The tight junctions in the healthy corneal epithelium provides a barrier against compounds larger than 1 nm. Topical administration of full-length immunoglobulins is generally ineffective because these drugs are too large to penetrate the intact cornea. In this study, bevacizumab was barely detected beyond the superficial layer the corneal epithelium in mice even after 7 days of topical application. The subconjunctivally injected bevacizumab in eyes with intact cornea penetrated well into the corneal stroma^23^.

In our IHC study, after topical application, the large bevacizumab was not detected in the retina. Lumen of the blood vessels in the inner retina and the optic nerve showed staining suggestive of systemic clearance. This was further supported by the western blot findings where signals were present in the plasma at 24 hours and 48 hours. No signals were detected in the eyes or in the brain. IHC staining of the outer segment of the photoreceptors for the small retinoschisin was present at 24 hours. The staining was not detected at 48 hours.

In a study, the pharmacokinetics of 1.25mg/0.05ml of Bevacizumab in the ocular tissue and the plasma of Pigmented rabbits was compared after administering the drug in the form of either repeated topical eye drops six times a day for 7 days, a single subconjunctival injection or a single intravitreal injection. Intravitreal injection was found to be the most effective route to retina/choroid and iris/ciliary body followed by the subconjunctival route. Topical administration showed poor distribution in the target tissues. Both intravitreal and subconjunctival routes resulted in high plasma concentrations^24^

In our study also, the intravitreal technique was more effective than the topical and subconjunctival techniques. Though the above-mentioned techniques target the posterior segment of the eye there is still no technique to effectively target the retina, the optic nerve and the visual pathway in the brain. A technique that can target both the brain and the eye can be useful in many age-related and neurodegenerative conditions. Therefore, a novel maxillofacial route was assessed in the mouse model.

The glymphatic pathway in the brain was first proposed in 2012. Based on experimental studies in mice, it was proposed that CSF enters the paravascular spaces in the penetrating arteries in the cortex in the same direction as the blood flow. CSF then passes through the AQP4 channels in the astrocytic endfeet surrounding the capillaries and enters the interstitial space. The ISF then enters the paravenous space and exits into the subarachnoid space, the venous sinuses, the perineural sheaths of the cranial and spinal nerves space and is transported to the cervical lymph nodes by the meningeal lymphatic channels^25, 26^.

In 2015, the existence of a glymphatic pathway in the visual pathway was proposed.^27, 28^ The optic nerve is surrounded by all the three meninges. CSF is present within the subarachnoid space. The central retinal artery pierces the dura and the arachnoid of the optic nerve adjacent to the globe and gives centrifugal branches during its course in the center of the optic nerve The retrobulbar region of the optic nerve is also supplied by pial vessels that send centripetal branches into the optic nerve. The optic nerve head and retrolaminar optic nerve are primarily drained by branches of the central retinal vein that courses through the center of the optic nerve alongside the central retinal artery. It drains into the superior and inferior ophthalmic veins which drain into the cavernous sinus, the pterygoid venous plexus and the facial vein.

In-vivo studies were conducted in mice to assess whether CSF enters the optic nerve via a glymphatic pathway. Fluorescent dextran tracers that were injected into the cisterna magna were found in the subarachnoid space surrounding the intra-orbital optic nerve. Tracers were also seen surrounding the blood vessels in the optic nerve in direct contact with the blood vessel endothelium as well as approximately 1-2 µm peripheral to the endothelium. Tracers were also seen around the blood vessels entering the optic nerve from the subarachnoid space. The tracers were located in the paravascular spaces that were lined externally by GFAP positive astrocytes.^29^

Recently, MRI studies were conducted to assess whether direct communication between the CSF and the extravascular compartment of the human visual pathway. Gadobutrol, a CSF tracer, was administered intrathecally and its distribution into the glymphatic paravascular pathways of the orbital and the intracranial segments of the optic nerve and the central visual pathway were studied. CSF tracer enrichment was seen in the optic nerve, optic tract and the primary visual cortex after 24 hours. The prechiasmatic cistern showed peak enhancement within 4 - 6 hours and the optic chiasma showed a peak within 6-9 hours. Tracer enhancement was also seen in the retrobulbar part of the optic nerve corresponding to the entrance of the central retinal artery^30^.

Another MRI study using intrathecal gadobutrol in human patients with idiopathic normal pressure hydrocephalus showed delayed enhancement and clearance of the tracer in the human visual pathway suggesting an impaired glymphatic pathway in the disease. The tracer distribution within the visual pathway depended on the availability of tracer in the CSF and seemed to be affected by the intracranial compliance ^31^.

Glymphatic pathway has been proposed to play a role in the clearance of β-amyloid in the brain and amyloid deposits in the retina. A new study assessed intraocular anterograde glymphatic clearance system in the eyes of mice by injecting HiLyte-594-tagged human β-amyloid into the vitreous and visualizing the tracer distribution one hour later. 3D reconstruction of light-sheet microscopy data showed that in addition to anterior routes, the tracer exited the eye along the optic nerve. Whole-mount preparation of the optic nerve confirmed the anterograde transport of the tracer along the nerve. It accumulated in the perivascular space of the veins rather than the arterioles^32^.

Sectioning and high-resolution imaging of the eye showed that, in addition to perivenous accumulation, the tracer was taken up by retinal ganglion cells, amacrine cells and transported along neuron-specific class III β**-**tubulin positive axons. The tracer accumulation peaked at 274 ± 20 μm from the optic nerve head and tapered off gradually in the distal nerve. The dural sheath surrounding the proximal segment of the optic nerve densely accumulated the tracer. This showed that after intravitreal delivery, the tracer is transported by axons and along the perivenous space in the optic nerve and drains into the dural lymph vessels and the cervical lymph nodes^32^.

Imaging after dual injections of hAβ tracer in the vitreous body in conjunction with a tracer in the cisterna magna highlighted tracer transport within the optic nerve in both anterograde and retrograde directions with limited spatial overlap. Tracers injected in the cisterna magna were transported along perivascular spaces of the arteries and the capillaries into the optic nerve Therefore in addition to the trabecular and uveoscleral outflow routes, the glymphatic pathway has also been proposed as a pathway for efflux of ocular fluid^32^.

It has been proposed that normal tension glaucoma could result from disturbance in the glymphatic pathway in the brain and high-tension glaucoma could result from disturbance in the ocular glymphatic system^33^.

Our study that was published earlier had used an oral and maxillofacial device to deliver drugs into the brain ^3^. In this in vivo mouse study, we used the novel maxillofacial drug delivery system to deliver drugs to the retina and optic nerve by targeting the glymphatic pathway. Drugs can be transported to the optic nerve from the brain through the glymphatic system and the CSF and be delivered into the retina.

In vivo studies have shown that human glymphatic system in the brain functions generally different from animals. Human studies have shown brain wide enrichment of CSF tracer and the enrichment occurs over a protracted period of over 24 hours. In animals the glymphatic is primarily cortical and the circulation is fast within 1-2 hours probably because of the smaller brain size^25^.

In this mouse study, the animals were sacrificed at 24 hours and 48 hours after the Avastin injection. IHC staining for bevacizumab was seen in the brain in the cortex, brainstem and cerebellum. Staining was stronger in the 24-hour tissue sections than in the 48-hour tissue sections. The walls of the parenchymal and the pial vessels were stained suggestive of clearance through the glymphatic pathway. The staining of the choroid plexus in spite of negligible drug in the plasma was suggestive of deeper penetration of the drug^34, 35^.

Our in vivo real-time imaging study in mice using the maxillofacial device that was published earlier showed delivery of a fluorescent dye (Indocyanine Green) into the brain. Though the drug was delivered to only one side of the maxillofacial region, it was noted that there was global distribution of the drug in the brain. But nasal drops applied topically on the nasal mucosa of the mouse showed distribution of the drug in the brain only at the midline. The study also showed that the drug was transported to the eyes. The maxillofacial route showed slightly more fluorescence in the contralateral eye^36^.

In this IHC study, 24 hours after the Avastin injection, staining was detected in the outer segment of the photoreceptors in the ipsilateral eye as well as the contralateral eye. Similarly staining was also detected in the ipsilateral optic nerve and the contralateral optic nerve. The staining was slightly more in the contralateral eye.

At 48 hours, staining decreased in the outer segment of the photoreceptor layer of both the ipsilateral and contralateral eye. The staining was still slightly more in the contralateral eye. Staining was present in the ipsilateral and contralateral optic nerve also. On injecting two doses of the drug, staining was very significant in the contralateral optic nerve at 48 hours suggestive of clearance of the drug through the glymphatic pathway of the optic nerve.

The contralateral enhancement in the optic nerve while targeting the glymphatic pathway can be explained by the neuroanatomy of the visual pathway. The visual impulses from the retina are carried through the optic nerve, the optic chiasma, the optic tract, the lateral geniculate ganglion and optic radiation to the visual cortex in the occipital lobe.

In all species, the majority of RGC axons project to the contralateral side at the optic chiasma. In primates, the number of ipsilateral and contralateral RGC axons is approximately equal^37^. In humans, ratio of crossed to uncrossed fibers in the chiasm of 53:47 ^38^. In mouse, only 3% to 5% of all RGC axons are ipsilateral^37^. In rodents, uncrossed axons also approach the chiasmal midline. In humans, the uncrossed axons are confined laterally from the optic nerve to the optic tract^39^.

A study that assessed anterograde diffusion of ICG was assessed after intravitreal injection in a rabbit model showed long term staining of the visual pathway. The ipsilateral optic nerve, the optic chiasma, the contralateral superior colliculus and the contralateral lateral geniculate nucleus exhibited ICG staining. Further, retrograde diffusion of ICG was assessed after microinjection into the lateral geniculate nucleus of rats. Fluorescence of the contralateral retinal ganglion cells and of the disc were observed^40^.

A study using a mouse model of laser induced ocular hypertension in which a retrograde tracer was applied on the surface of the superior colliculus one week prior to animal processing showed retrograde transport of the tracer to the retinal ganglion cells^41^.

Injections of fluorescent retrograde tracers into the lateral geniculate ganglion and the superior colliculus of albino rats resulted in predominant staining of the contralateral retinal ganglion cells^42^.

In studies involving experimental glaucoma in mice, the astroglia and Mueller cells in the contralateral eye showed reactive signs that differed from the experimental eye and the naïve eye. Similarly, the microglia in the contralateral eye showed up regulation of MHC-II^43^.

The above findings show that the distribution of the drug into the contralateral retina and optic nerve in the rodent model could be because of the significant crossover of axons to the contralateral side at the optic chiasma. The drug from the brain could have also distributed into the optic nerve through the glymphatic pathway.

Our western blot analysis at 24 hours after administration of Avastin in the right maxillary sinus region using the maxillofacial technique showed that signals were present in both the right and left halves of the brain. The right eye and the left eye showed signals. IHC studies showed mild enhancement in the staining of the contralateral eye when compared to the ipsilateral eye. This finding supports that the drug is transported from the brain to the contralateral optic nerve and the contralateral retina. The persistence of staining in the optic nerve at 48 hours after staining in the photoreceptor layer of the retina has disappeared further shows that the optic nerve could also provide an elimination route. Therefore, the glymphatic pathway can play a significant role in ocular drug delivery.

Further, our western blot analysis at 24 hours after administration of Avastin through the maxillofacial route showed positive signals in the plasma along with positive signals in the right and left eye.

Following intravitreal injection into the eye, bevacizumab has been also detected in the fellow non-injected eye in many studies. Preclinical studies in rabbits also have shown low concentration of bevacizumab in the fellow non-injected eye. Following intravitreal injection of the drug into the experimental eye, a maximum concentration in the serum was detected 8 days after the injection and declined by 29 days. The vitreous of the contralateral non-injected eye showed a gradual increase in concentration from day 1 to 4 weeks. The concentration in aqueous peaked at one week and declined at 4 weeks. The concentration in the aqueous humor was higher than the vitreous at all time points before 21 days. Hence, entry from the systemic circulation into the eye through the anterior route was proposed.^14^.

In the rabbit study that compared topical route, subconjunctival route and the intravitreal route for administering Bevacizumab, the anterior structures of the fellow eye namely the iris/ ciliary had more drug concentration than the retina/choroid in all the three routes and the vitreous had the least concentration. Distribution into the fellow eye was proposed to be due to systemic circulation^24^.

Clinical studies have also reported that unilateral bevacizumab injections in patients with bilateral diabetic macular edema is often associated with a bilateral response. This effect has been proposed to be due to escape of bevacizumab into the systemic circulation that may be facilitated by the breakdown of the blood retinal barrier in eyes with diabetic macular edema^44^.

In the rabbit studies cited above, the aqueous and the anterior structures showed higher concentration of the drug than the posterior segment. The drug was detected in the contralateral eye as early as day 1. The aqueous concentration was higher than the vitreous at all times. It has been shown that larger molecules in the vitreous tend to be cleared through the anterior routes. Therefore, it is possible that the higher concentration of bevacizumab in the aqueous is because of the anterior clearance from the vitreous.

Our IHC study showed staining of the optic nerve at 24 hours and 48 hours. In studies involving experimental glaucoma in mice, the astroglia and Mueller cells in the contralateral eye showed reactive signs that differed from the experimental eye and the naïve eye^24^. If these small quantities of cytokines produced in the experimental eye need to be initially transported through the systemic circulation to reach the cells in the contralateral eye, systemic degradation of the cytokines is more likely to happen. This should have led to loss of signaling at the contralateral target tissues.

Similarly, large drugs like bevacizumab cannot cross the intact blood retinal barrier. Systemic route may be a plausible route only if the blood retinal barrier is either compromised as in bilateral diabetic macular edema or if the systemic concentration of the drug is high. Therefore, a local transport using the glymphatic pathway at the optic chiasma is more plausible.

## CONCLUSION

1. In this study, the maxillofacial technique and the intravitreal injection effectively delivered bevacizumab to the retina. The maxillofacial technique could also target the optic nerve
2. The glymphatic pathways of the brain and the optic nerve can be targeted for drug delivery using the maxillofacial technique. This can be useful for conditions like optic neuritis, multiple sclerosis and other neurodegenerative conditions involving the brain and the eye.
3. The maxillofacial technique can deliver drugs into the brain and both the eyes. Early enhancement of the tracers in the pre-chiasmatic cistern and optic chiasma have been reported in MRI studies using intrathecal tracers. In this study, early distribution of the drugs into the optic chiasma from the brain via the glymphatic system could have resulted in the delivery of the drugs into both the eyes through the respective optic nerves.
4. In this study, contralateral optic nerve and retina showed slightly more staining in the maxillofacial technique. The distribution of the drug in the CSF at the optic chiasma, the crossover of the axons of the optic nerve at the optic chiasma and the delivery of the drug to the contralateral eye through the glymphatic system of the optic nerve could have contributed to the observed staining in the contralateral eye.
5. Large drugs like bevacizumab do not cross the blood retinal barriers. Therefore, systemic transport of the drug from the ipsilateral eye to the contralateral eye is possible only in disease conditions wherein the blood retinal barriers are damaged. In healthy eyes with intact blood retinal barriers, it is more likely that a redistribution of the drug may occur in the CSF at the optic chiasma and the drug may be transported to the contralateral eye through the ocular glymphatic pathways.

## AUTHOR’S CONTRIBUTION

The new concept to explain the presence of drug in the non-injected contralateral eye based on the glymphatic pathway and the crossover of the optic nerve fibers at the optic chiasma was developed by Jayamuruga Pandian Arunachalam, Subbulakshmi Chidambaram, Anoop U.R, Kavita Verma and Rahini Rajendran. All the authors contributed to the study.

## FUNDING

This study is supported financially by the Science & Engineering Research Board (SERB), Department of Science & Technology (DST), Government of India (EMR/2017/005413).

## CONFLICT OF INTERESTS

The authors, Anoop. U. R and Kavita Verma are stated as the inventors and applicants in the granted patents US11,207,461B2 and AU2016300184; and in the patent application PCT/IB2016/053899 with National Phase Entry into India European Patent Office and Canada.

## ETHICS COMMITTEE APPROVAL

All the animal experiments of the study are approved by the Institutional Animal Ethics Committee, Sri Balaji Vidyapeeth [Deemed to be University] (03/IAEC/MG/11/2018-II).

## ACKNOWLEDGMENT

We thank DST SERB (Government of India) for providing funding to carry out the study. We are grateful to the Central Inter-Disciplinary Research Facility and Sri Balaji Vidyapeeth (Deemed to be University) for providing the necessary facilities to conduct the study.

We would like to thank Dr.R.Pachaiappan SRM Institute of Science and Technology, Kattankulathur, Chengalpattu District, Tamil Nadu 603203, India for the LCMS/MS study. We would like to acknowledge Dr. John Baliah, Department of Oral Medicine and Radiology, Indira Gandhi Institute of Dental Sciences and Research, Sri Balaji Vidyapeeth (Deemed to be University) for radiovisiography radiographs.

---

**Figure 1:**
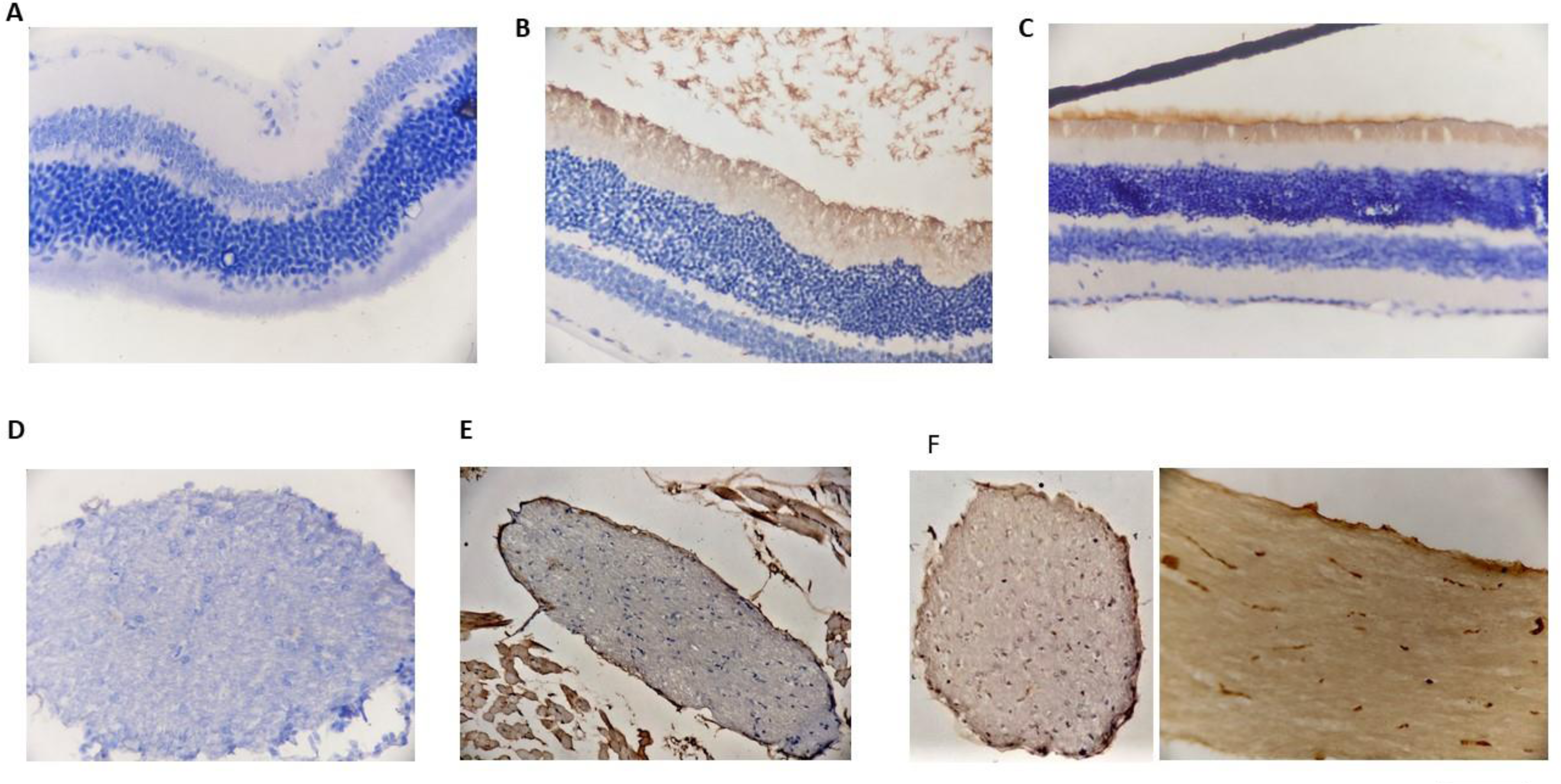
**A**-shows the control retina; **B** - shows the retina after intravitreal injection, **C** - shows the retina after maxillofacial administration, **D -** shows the control optic nerve, **E -** shows the optic nerve after intravitreal injection, **F -** shows the optic nerve after maxillofacial administration.

**Figure 2:**
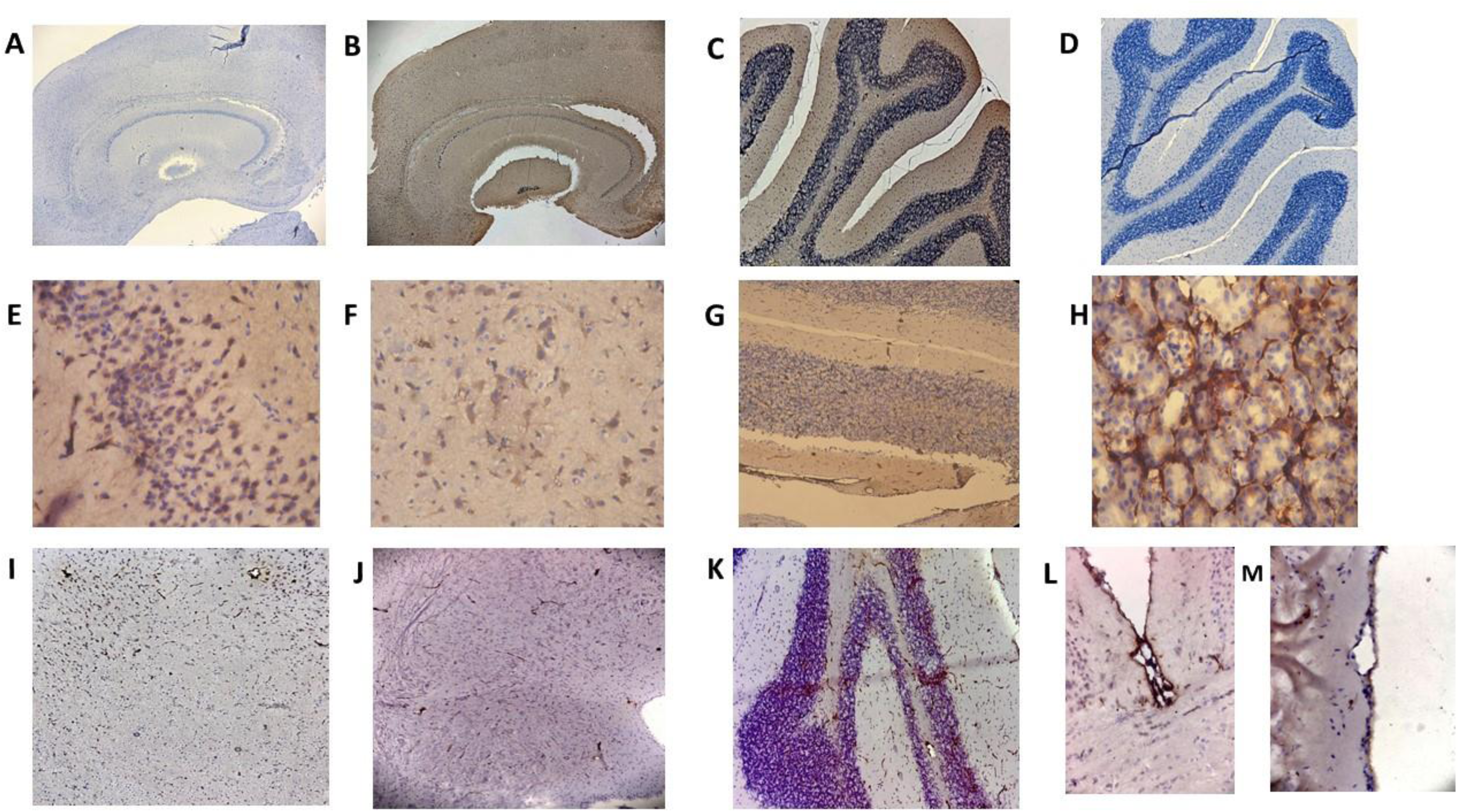
**A**-shows the negative control brain; **B** - shows the positive control brain; **C** - shows the positive control cerebellum **D -** shows the negative control cerebellum **E -** shows the cortex at 24 hours after administration of Avastin through maxillofacial route, **F -** shows the Pons 24 hours after administration of Avastin through maxillofacial route, **G -** shows the cerebellum 24 hours after administration of Avastin through maxillofacial route, **F -** shows the kidney 24 hours after administration of Avastin through maxillofacial route, **I -** shows the cortex 48 hours after administration of Avastin through maxillofacial route. **J -** shows the Pons 48 hours after administration of Avastin through maxillofacial route, **K -** shows the cerebellum 48 hours after administration of Avastin through maxillofacial route, **L, M** show the pial vessels with staining in the walls 48 hours after administration of Avastin through maxillofacial route.

**Figure 3:**
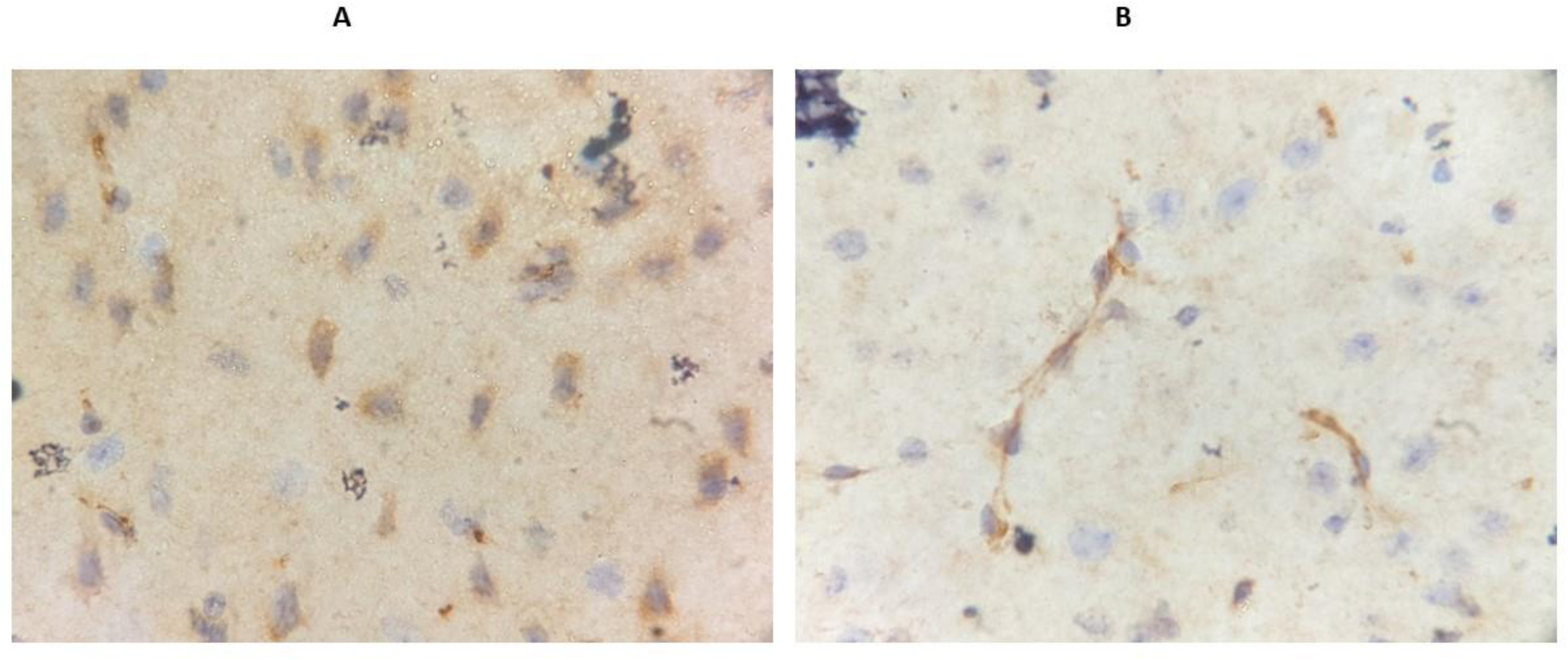
**A**-Shows brain 24 hours after administration of RS1-FLAG through maxillofacial route, B- Shows brain 48 hours after administration of RS1-FLAG through maxillofacial route

